# The histone demethylase Kdm5 and the ARGONAUTE proteins Piwi and Aubergine regulate female abdominal pigmentation in *Drosophila melanogaster*

**DOI:** 10.64898/2026.08.27.747613

**Authors:** Raphael Narbey, Sandra De Castro, Sophie Louvet-Vallée, Michel Gho, Frédérique Peronnet, Jean-Michel Gibert, Emmanuèle Mouchel-Vielh

## Abstract

Insect pigmentation is an ecologically critical trait influencing many physiological processes. In *Drosophila melanogaster*, abdominal pigmentation is sexually dimorphic: males have fully pigmented posterior segments, while females exhibit a posterior melanin stripe. Pigmentation relies on the expression of pigmentation genes that encode enzymes involved in pigment synthesis. These genes are tightly regulated during pupal and young adult stages. To expand the gene regulatory network of pigmentation genes, we conducted an RNAi screen using the *yellow-Gal4* driver, expressed during the pupal stage in abdominal epidermis. One of the candidates from this screen, *Kdm5*, encodes a histone demethylase erasing the H3K4me3 histone mark catalyzed by the histone methyl-transferase Trithorax (Trx). We show that *Kdm5* down-regulation reduces abdominal pigmentation, mimicking *trx* down-regulation. Kdm5 activates melanin production through regulation of the pigmentation gene *tan*. Transcriptomic analyses reveal that Kdm5 and Trx share many targets in pupal abdominal epidermis, including piRNA pathway components such as *piwi* and *aubergine*. These piRNA components, originally associated with transposon silencing in the germline, also function in some somatic tissues such as the nervous system, the fat body or the gut. We demonstrate that Piwi and Aubergine participate in female abdominal pigmentation establishment, without evident piRNA production. We also show that Kdm5 and Piwi act not only in pupal abdominal epidermis but also in pupal fat body. This study therefore expands the regulatory network of pigmentation genes. It identifies a new somatic function for Kdm5 and Piwi and reveals a role for pupal fat body in female abdominal pigmentation regulation.

**Article summary:** We investigated Drosophila female abdominal pigmentation to enrich the regulatory network of pigmentation genes. Using a Gal4 driver expressed in abdominal epidermis, we conducted an RNAi screen. We discovered that the histone demethylase Kdm5 activates the pigmentation gene *tan*. Using transcriptomic analysis and genetics experiments, we showed that the piRNA genes *piwi* and *aub* act downstream of Kdm5 to regulate *tan* expression, without evident piRNA production. Moreover, *Kdm5* and *piwi* act not only in the pupal epidermis but also in the pupal fat body, thus showing for the first time a role of this tissue in abdominal pigmentation regulation.

## Introduction

Insect pigmentation is an ecologically relevant trait linked to many adaptive processes such as pheromone production, thermoregulation, resistance to UV, desiccation, pathogen or parasites (reviewed in [1]). In many species of *Drosophilidae*, abdominal pigmentation is characterized by sexual dimorphism. In *Drosophila melanogaster* males, the posterior most segments are fully pigmented. In contrast in females, posterior segment pigmentation is restricted to a posterior stripe, similar to that present on the anterior segments in both sexes. Abdominal pigmentation relies on the synthesis of several pigments whose precursors are produced by epidermal cells and secreted into the developing cuticle at the late pupal stage. The enzymes involved in the pigment synthesis pathway are encoded by structural genes, called thereafter pigmentation genes, expressed from the second half of the pupal life to the beginning of the adulthood [2].

In *D. melanogaster* and other *Drosophila* species, several studies have revealed that the spatial and sex-specific patterning of abdominal pigmentation is tightly controlled by transcription factors involved in diverse developmental processes, which directly or indirectly regulate pigmentation gene transcription [1]. For example, the Hox proteins Abdominal-A and Abdominal-B regulate the pigmentation genes *tan* (*t*) and *yellow* (*y*) [3,4]. The transcriptional repressors Bric-a-Brac 1 and 2, which are encoded by the two paralogous genes *bab1* and *bab2*, repress *y* and *t*, thus negatively controlling melanin production [5–8]. To enrich the gene regulatory network of pigmentation genes in *D. melanogaster*, several genetic screens have been performed, based on RNAi and CRISPR/Cas9 technologies [9–11]. The Gal4 driver used in these screens was the early *pannier-Gal4* driver (*pnGal4*) expressed throughout development, from the embryo to the adult [12]. In these screens, only genes encoding transcription factors were tested.

In this study, we describe an RNAi genetic screen using the late *yellow-Gal4* (*yGal4*) driver expressed during the pupal life in the abdominal epidermis [3]. Using this driver helped to prevent defects due to early inactivation of developmental genes. Our long-standing interest in female abdominal thermal plasticity led us to focus on females only [8,13]. Moreover, we included in our screen not only genes encoding transcription factors, but also genes encoding chromatin regulators. Indeed, as illustrated by previous studies, regulation of chromatin structure plays a role in the establishment of abdominal pigmentation. For example, the histone-methyl transferase Grappa (Gpp), which catalyzes the active histone mark H3K79me3, regulates different pigmentation genes depending on the developmental stage: Gpp has an early role in larvae leading to *y* repression and a latter role in pupae leading to *t* activation [14]. Trithorax (Trx), another histone-methyl transferase which catalyzes the active mark H3K4me3, is also involved in abdominal pigmentation, activating *t* and repressing another pigmentation gene, *ebony* (*e*) [13,15].

Among candidate genes from this screen, we focused on *Kdm5*, also called *little imaginal disc* (*lid*). *Kdm5* down-regulation induced a decrease of abdominal pigmentation, producing a phenotype similar to that induced by *trx* down-regulation. First identified as a gene from the Trithorax-Group [16], *Kdm5* encodes a histone demethylase erasing the H3K4me3 mark [17–19]. In *D. melanogaster*, Kdm5 is one of the 12 histone lysine demethylases with a catalytic JumonjiC (JmjC) demethylase domain [20]. Kdm5 is essential for viability, although its demethylase activity is dispensable to ensure a proper development [21,22]. This observation highlights the importance of the other protein domains and regulatory activities independently of H3K4me3 demethylation. During oogenesis, Kdm5 is a major regulator of the oocyte epigenome [23]. It activates the expression of *deadhead* (*dhd*), a critical effector of the oocyte-to-zygote transition, making it a key actor of early development [24]. It also plays a role in neuronal function, synaptic morphology and neurotransmission, through demethylase-dependent and independent mechanisms [25,26]. Kdm5 was shown to physically interact with various chromatin regulators and transcription factors, such as the SIN3A histone deacetylase complex [27,28], the Myc oncoprotein [29] or the transcription factor Foxo [30]. Other partners of Kdm5 were recently identified using the TurboID-mediated proximity technology, including members of the SWI/SNF, NURF, NSL and Mediator complexes [31]. Contrary to what one might expect from its enzymatic activity in erasing the active mark H3K4me3, Kdm5 acts mainly as a transcriptional co-activator. Indeed, it antagonizes heterochromatin-dependent gene silencing and is required to maintain the pattern of H3 acetylation, an active histone mark [32]. Kdm5 preferentially binds the transcription start site of actively transcribed genes where it co-localizes with H3K4me3. Consistently, the transcription of genes bound by Kdm5 decreases upon Kdm5 depletion [33,34]. It was proposed that transient demethylation of the promoters by Kdm5 could reset chromatin to an unmethylated state at the end of a transcription cycle, thereby allowing a new cycle to begin [34].

In this study, we demonstrate that Kdm5 activates melanin production in the female abdominal epidermis through activation of the pigmentation gene *t* and repression of one of its regulators, *bab1*. RNA sequencing experiments reveal that Kdm5 and Trx share many common transcriptional targets in the pupal abdominal epidermis, including several actors of the piRNA metabolism. piRNA are small non-coding RNA present in germline cells and gonadal tissues of many organisms where they silence transposable elements, thus ensuring genome integrity. They arise from the transcription of specific loci, called piRNA clusters, composed of fragments of transposable elements. Many proteins participate in the piRNA pathway, including several ARGONAUTE proteins of the PIWI family, such as Ago3, Piwi and Aubergine (Aub) in drosophila (for a review, see [35]). Small RNA that display the features of piRNA have also been identified in non-gonadic somatic tissues of diverse species, suggesting that the piRNA pathway or some of its actors individually could also act outside the gonads [36]. For example, a function of piRNA and piRNA pathway proteins has been described in the nervous system of several organisms, such as *C. elegans*, drosophila, aplysia, mouse (for a review, see [37]). A functional piRNA pathway also exists in the adult drosophila fat body, where it ensures metabolic homeostasis [38]. In the gut of drosophila adults, Piwi is expressed in the intestinal stem cells where it participates in the maintenance of homeostasis by transposon silencing [39,40]. In addition to Piwi, Aub is also required to regulate proliferation of adult intestinal stem cells in drosophila [41]. Apart from their role in the repression of transposable elements, other cellular functions of Piwi and Aub have been described in drosophila and mammals. Some of them seem to be independent of piRNA production (reviewed in [42]). We thus wondered whether the piRNA pathway or some of its actors participates in the establishment of female abdominal pigmentation. Using genetic and molecular experiments, we demonstrate that the ARGONAUTE proteins Piwi and Aub, whose expression is activated by Kdm5, act in female abdominal epidermis, without evident piRNA production, to regulate melanin production through the control of *tan* expression. Furthermore, in addition to their role in abdominal epidermis, Kdm5 and Piwi also act in the pupal fat body to regulate abdominal pigmentation. Our study therefore reveals a new somatic role for Piwi and demonstrates for the first time a role of the pupal fat body in the regulation of abdominal pigmentation.

## Material and methods

### Fly stocks and genetic crosses

All crosses were performed at 25°C, using 10 females and 6 males. For control crosses, a *w^1118^* line was used. The lines used in the experiments, listed in Supplementary Table 1, were mainly obtained from the Bloomington Stock Center except a few of them. The *yGal4* transgene was introgressed into the *w^1118^* background for 10 generations. In all experiments except that of Supplementary Figure 2, the *RNAi-Kdm5* line used was BDSC_28944. The *5’tan-GFP* line was generated as previously described [8]. The *RNAi* lines tested in the genetic screen were from the *TRIP RNAi* collection [44] and listed in Supplementary Table 2. In the screen, for each tested *RNAi* line, we compared by eye the abdominal pigmentation of the female progeny from *yGal4* females crossed with males of the tested *RNAi* line with that of two control crosses: (1) *yGal4* females crossed with either *RNAi-GFP-attP2* or *RNAi-GFP-attP40* males, depending on the insertion platform of the *RNAi* transgene; (2) *w^1118^* females crossed with males of the tested *RNAi* line. Only the *RNAi* lines giving congruent results with the two controls are listed in Supplementary Table 2.

### Preparation of cuticles, image acquisition and quantification

For pigmentation analyses, adult females between 3 and 5 days old were stored in 70% ethanol for ten days. The abdominal cuticles were cut just beyond the dorsal midline and dehydrated in 100% ethanol for 5 minutes. After dehydration, the cuticles were mounted in Euparal (Roth) and imaged with a Leica DC480 digital camera using the Leica IM50 Image Manager software. For GFP analyses, the cuticles and the underlying epidermes of adult abdomens were dissected in PBS, fixed 20 minutes in 3.7% paraformaldehyde in PBS, washed twice 10 minutes in PBS, mounted in Mowiol and imaged with a micro-apotome (Zeiss). Adult pigmentation and GFP intensity quantifications were performed as previously described [8,13].

### Analysis of the *UAS-H2B-YFP* reporter transgene expression in larval and pupal tissues

*Lsp-2Gal4* or *yGal4* females were crossed with males containing a *UAS-H2B-YFP* transgene [66]. Third instar larvae were dissected according to the method described in [67]. Briefly, larvae were immobilized at both ends with pins on a Sylgar® surface and opened along the dorsal line with micro-scissors. The lateral epithelium was then opened and held in place with pins to expose the internal organs. A similar experimental strategy was applied to pharates, except that the animal was immobilized with two pins in the cephalic region. Both types of specimens were fixed in 4% PFA for 20 minutes, washed three times with PBT, and mounted for microscopic observation in a 1:1 PBS/glycerol solution. Tissues were observed using an Olympus BX41 fluorescence microscope equipped with a Yokogawa spinning disk and a Retiga R1 CCD camera controlled by Metaview software (Universal Imaging). Images were obtained by maximum projection of a Z stack composed of 2 to 6 optical sections (spaced 1µm apart).

### RT-qPCR experiments

The RNeasy Mini kit (Qiagen) was used to extract RNA from female dissected tissues: (1) posterior abdominal epidermis of 30 old female pupae (stage P12(i) defined by Bainbridge and Bownes, [68]); (2) posterior abdominal epidermis of 30 young female adults within two hours after eclosion; (3) pupal fat body of 20 P12(i) old female pupae. After treatment of RNA with DNAse1, cDNAs were synthesized with the LunaScript RT SuperMix kit (New England Biolabs) using random primers. RT-qPCR experiments were performed in a CFX96 system using SsoFast EvaGreen SuperMix (Biorad). Expression was quantified following the Pfaffl method using the geometric mean of the expression of *RP49* and *Spt6* reference genes for normalization [69,70]. The primers used in these experiments are listed in Supplementary Table 3.

### RNA sequencing and bioinformatic analyses

Total RNA from pools of 30 posterior abdominal epidermis from P12(i) old female pupae were extracted using the RNeasy Mini Kit (Qiagen) and treated with DNAse1. Libraries preparation (Illumina TruSeq Stranded Total-RNA Ribozero Gold kit) and sequencing (Illumina NovaSeq 600) were carried out by the FASTERIS company, with three biological replicates *per* genotype. Reads were aligned with the *D. melanogaster* genome (dm6, r6.41) using HISAT2 (Galaxy v2.1.0) and counted using FeatureCounts (Galaxy v1.6.4). Differential expression analysis was performed with DEseq2 (Galaxy v1.6.0.6). Supplementary Table 4 presents the total number of reads and the reads used for differential analysis. Genes with an adjusted p-value below 0.05 were considered as differentially expressed. Gene Ontology analysis was performed using DAVID [71]. The RNAseq data are available in the GEO database (https://www.ncbi.nlm.nih.gov/geo/info/seq.html) under accession number GSE334852.

### Small RNA sequencing and bioinformatic analyses

RNA from the posterior abdominal epidermis of 30 P(12)i old female pupae or from the pupal fat body of 20 P(12)i old female pupae were extracted with the TRIzol reagent (Thermo Fisher Scientific) and treated with DNAse1. Libraries preparation (UMI small RNA kit), sequencing (DNBSEQ) and filtering of raw reads (SOAPnuke package) were carried out by the BGI company (one replicate *per* genotype). sR_Bowtie (Galaxy v2.3.0) was used to align reads on a version of the *D. melanogaster* genome release 6 in which the unmapped reads, the Y, and the mitochondrial chromosomes were removed (GEO accession number GSE203279, [52]). For comparison between libraries, read counts were normalized to one million (rpm). Successive sR_Bowtie alignments were performed on the total aligned reads to remove mi-RNA, t-RNA and misc-RNA sequences (sequences downloaded from FlyBase). The size profiling of the remaining reads, further called cleaned reads, was analyzed using Small RNA Map (Galaxy v3.1.1). The number of reads for each size category (21-nt, 22-nt, 23-29-nt) was counted with Clip Adapter (Galaxy v2.5.1) and Count Alignments (Galaxy v4.1.1). The 23-29-nt reads containing a uracil in first position were counted using the Line/Word/Character Count command (Galaxy v1.0.0). Logos were generated using Sequence Logo (Galaxy v3.5.0) after trimming of the reads to 15-nt (Trim Sequences, Galaxy v1.0.2). Supplementary Tables 5 and 6 present the characteristics of the small RNA libraries. The small RNAseq data from the pupal abdominal epidermis and fat body are available in the GEO database (https://www.ncbi.nlm.nih.gov/geo/info/seq.html) under accession numbers GSE334956 and GSE334957, respectively.

## Results

### *Kdm5*, encoding the histone demethylase 5, regulates female abdominal pigmentation

To identify new transcription factors and chromatin regulators involved in the abdominal pigmentation of female *D. melanogaster*, we performed an RNAi screen using the *yellow-Gal4* (*yGal4*) driver. This driver is expressed in the abdominal epidermis during the late pupal stage [3]. The list of genes tested in this screen was established using the FlyTF database [43] (categories “putative transcription factors” and “protein involved in putative chromatin processes”). After comparing this list with preliminary RNA-seq data from the abdominal epidermis of wild type female pupae, we selected 494 genes expressed in this tissue, and for which a line from the *TRIP* RNAi collection was available [44]. For each tested line, the abdominal pigmentation of *yGal4>RNAi* females was compared to that of two control genotypes: *yGal4>RNAi-GFP* females and heterozygous *RNAi* females. The latter were obtained by crossing the tested RNAi line with control *w^1118^* flies since the *yGal4* driver was introgressed into the *w^1118^* background. The 476 genes for which congruent results were obtained with the two controls are listed in Supplementary Table 2. Most of the genes (n=432) induced no change in pigmentation, whereas 3 genes induced lethality. A decrease or increase of female abdominal pigmentation was observed for 36 genes and 5 genes, respectively. Thirty of these 41 genes were previously tested in an RNAi screen performed using the *pannier-Gal4* (*pnGal4*) driver expressed earlier during development, and 7 of them were found to induce a phenotype [9] (Supplementary Table 2).

Among genes inducing a pigmentation decrease with the *yGal4* driver, we were particularly interested in *Kdm5*. *Kdm5*, initially named *little imaginal discs* (*lid*), was first identified in a genetic screen for new Trithorax-group genes [16]. It encodes the histone demethylase 5 that demethyltes the histone mark H3K4me3 [17–19]. The methylation of H3K4 can be catalyzed by the histone methyl-transferases Trithorax (Trx), Trithorax-related (Trr) and Set1 [45]. Interestingly, we have shown previously that RNAi-mediated down-regulation of *trx* using the *yGal4* driver also induces a decrease in female abdominal pigmentation [13]. To confirm the results of the *yGal4* screen, we compared pigmentation intensity of the A5, A6 and A7 segments between control females (homozygous or heterozygous for the *yGal4* driver) and heterozygous or homozygous *yGal4>RNAi-Kdm5* females (*RNAi-Kdm5* line: BDSC_28944). We also quantified pigmentation of homozygous *yGal4>UAS-Kdm5* females to test the effect on pigmentation of *Kdm5* over-expression (Figure 1). The efficiencies of the *RNAi-Kdm5* and *UAS-Kdm5* transgenes were checked by RT-qPCR quantification of *Kdm5* expression (Supplementary Figure 1). Compared to *yGal4* control flies, *Kdm5* down-regulation induced a significant decrease of pigmentation in A6 for the heterozygous females and in A5, A6 and A7 for the homozygous ones. On the opposite, A5, A6 and A7 pigmentation was increased by *Kdm5* over-expression (Figure 1). A decrease in pigmentation was also observed when comparing *yGal4>RNAi-Kdm5* with *RNAi-Kdm5* heterozygous females or when using another *RNAi-Kdm5* line (BDSC_36652) (Supplementary Figure 2). Furthermore, a pigmentation decrease was also induced when using the *pnGal4* driver to down-regulate *Kdm5* with the two RNAi lines (Supplementary Figure 2). In conclusion, these results indicate that *Kdm5* participates in the establishment of female abdominal pigmentation and positively regulates melanin production.

**Figure 1.**
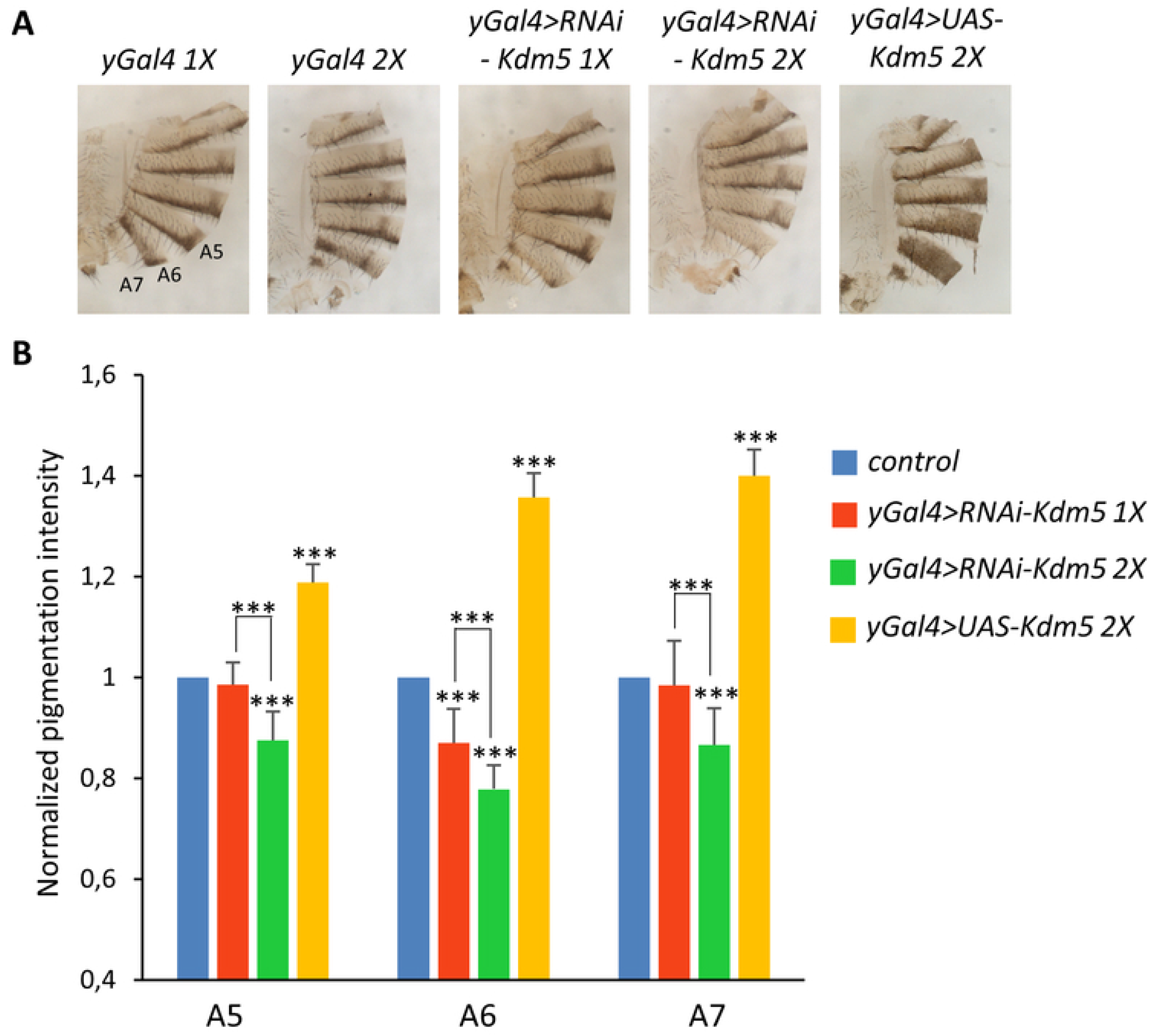
*Kdm5* positively regulates melanin production in abdominal epidermis of female drosophila. A. Abdominal cuticles of females heterozygous (1X) or homozygous (2X) of the following genotypes: control females (*yGal4*), females expressing the *RNAi-Kdm5* transgene (*yGal4>RNAi-Kdm5*) or the *UAS-Kdm5* transgene (*yGal4>UAS-Kdm5*). Cuticles were cut beyond the dorsal midline. A5, A6, A7: posterior abdominal segments. B. Quantification of A5, A6 and A7 pigmentation intensity in control *yGal4*, *yGal4>RNAi-Kdm5* and *yGal4>UAS-Kdm5* females (n=15 *per* genotype, t-test; ***: p<0.001). Pigmentation intensity of control flies was normalized to 1. Heterozygous *yGal4>RNAi-Kdm5* females were compared to heterozygous *yGal4* females, whereas homozygous *yGal4>RNAi-Kdm5* and *yGal4>UAS-Kdm5* females were compared to homozygous *yGal4* females. Down-regulation of *Kdm5* induced a significant decrease of pigmentation, whereas its over-expression increased pigmentation. Homozygous *yGal4>RNAi-Kdm5* flies were significantly lighter than heterozygous ones, suggesting a dose effect.

### Kdm5 activates the expression of the pigmentation gene *tan* through regulation of the transcription factor Bab1

In drosophila, the synthesis pathway of cuticular abdominal pigments involves several enzymes, encoded by pigmentation genes. These genes are sequentially expressed in the abdominal epidermis, from the second half of the pupal life to the beginning of adulthood [2,46]. We have previously shown that one of these genes, *tan*, which encodes a hydrolase involved in melanin production [47], was activated by the histone methyl-transferases Trx and Grappa (Gpp), catalyzing the active histone marks H3K4me3 and H3K79me3, respectively [13,14]. Furthermore, *tan* is the major effector of thermal plasticity of abdominal pigmentation. Indeed, temperature modulates the expression of *tan* through the activity of its abdominal enhancer, *t-MSE* [3,13]. As *Kdm5* is involved in abdominal pigmentation establishment, we sought to identify the pigmentation genes it regulates. We thus performed RT-qPCR experiments to quantify the expression of pigmentation genes in the posterior epidermis (A5, A6 and A7 segments) of young control female adults (*yGal4*) or female adults with a deregulation of *Kdm5* (homozygous *yGal4>RNAi-Kdm5* or *yGal4>UAS-Kdm5* females) (Figure 2A). *Kdm5* down-regulation induced a strong decrease of *tan* expression (fold-change: 0.09, p<0.001). Conversely, *tan* was up-regulated in females with *Kdm5* over-expression (fold-change: 2.3, p<0.001). The expression of two other pigmentation genes, *DDC* and *black*, was also slightly modified. The transcriptional regulation of *tan* by Kdm5 was confirmed by measuring the expression of the *5’tan-GFP* reporter transgene in abdominal epidermis of young females (Supplementary Figure 3). In this transgene, the expression of a nuclear GFP is controlled by *tan* 5’-regulatory sequences contained in a 4kb-sequence upstream of the transcription start site of *tan* and including the *t-MSE* abdominal enhancer. *Kdm5* down-regulation induced a decrease of *GFP* expression in A6 and A7 segments (Supplementary Figure 3A-B), whereas *GFP* expression was increased by *Kdm5* over-expression (Supplementary Figure 3C-D). Interestingly, no modification of *GFP* expression was observed when using a *t-MSE-GFP* reporter transgene (Supplementary Figure 3E-F), showing that the *t-MSE* enhancer alone was not sufficient to mediate the effect of Kdm5 on *tan* expression.

**Figure 2.**
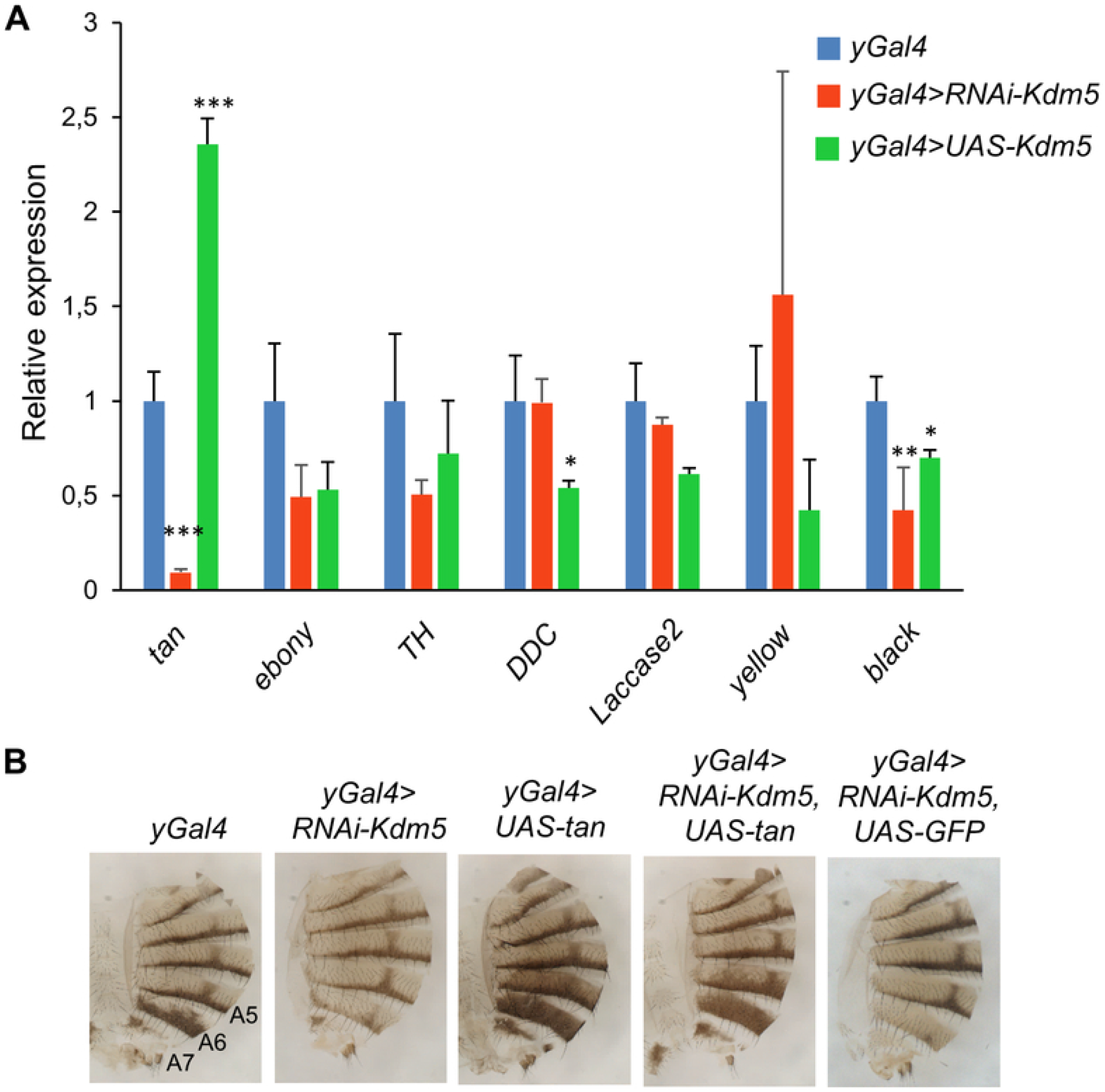
Kdm5 activates the expression of the pigmentation gene *tan*. A. Quantification of the expression of pigmentation genes in posterior abdominal epidermes (A5, A6, A7 segments) of homozygous young females either control (*yGal4*), or with down-regulation or over-expression of *Kdm5* (*yGal4>RNAi-Kdm5* and *yGal4>UAS-Kdm5*, respectively). The expression levels in *yGal4>RNAi-Kdm5* and *yGal4>UAS-Kdm5* flies were normalized to the expression levels in *yGal4* flies. The level of *tan* expression was significantly lowered by *Kdm5* down-regulation and increased by *Kdm5* over-expression (pools of 30 epidermes, n=3, normalization using the geometric mean of *RP49* and *Spt6* reference genes, t-test: *: p<0.05; **: p<0.01; ***: p<0.001). B. Abdominal cuticles of heterozygous females either control (*yGal4*), expressing an *RNAi-Kdm5* transgene (*yGal4>RNAi-Kdm5*), a *UAS-tan* transgene (*yGal4>UAS-tan*), these two transgenes together (*yGal4>RNAi-Kdm5, UAS-tan*) or the *RNAi-Kdm5* transgene and a *UAS-GFP* transgene (*yGal4>RNAi-Kdm5, UAS-GFP*). The pigmentation decrease induced by *Kdm5* down-regulation was suppressed by *tan* over-expression. This suppression was not due to Gal4 titration since no change in pigmentation was observed in *yGal4>RNAi-Kdm5, UAS-GFP* flies compared to *yGal4>RNAi-Kdm5* flies.

To confirm the regulation of *tan* by Kdm5, we analyzed pigmentation of flies with *tan* over-expression and *Kdm5* down-regulation (Figure 2B). As expected, females with *tan* over-expression (*yGal4>UAS-tan*) were much more pigmented than control flies (*yGal4*), whereas it was the opposite for females with *Kdm5* down-regulation (*yGal4>RNAi-Kdm5*). In *yGal4>RNAi-Kdm5, UAS-tan* females, *tan* over-expression suppressed the pigmentation decrease induced by *Kdm5* down-regulation, thus confirming that the decrease of *tan* expression was responsible for the depigmentation phenotype induced by Kdm5 depletion.

In a previous study, we have shown that, in pupal female abdominal epidermis, *tan* was repressed by the Bab1 and Bab2 transcription factors [8]. Bab1 and Bab2 are encoded by two paralogue genes, *bab1* and *bab2* (collectively named *bab*), that act during the pupal stage to negatively control melanin production in abdominal epidermis [5,6]. We therefore wondered whether the regulation of *tan* by Kdm5 was mediated by *bab*. Quantification of *bab* expression in pupal epidermes of control females (*yGal4*) or females with *Kdm5* down-regulation (*yGal4>RNAi-Kdm5*) revealed that *bab1*, but not *bab2*, was repressed by Kdm5 (Figure 3A). Furthermore, *bab1* was epistatic on *Kdm5*, as females that down-regulated both *Kdm5* and *bab1* exhibited the same increase of pigmentation as females with a down-regulation of *bab1* only (Figure 3B). Thus, *bab1* acts downstream of *Kdm5* within the gene network that controls *tan* expression.

**Figure 3.**
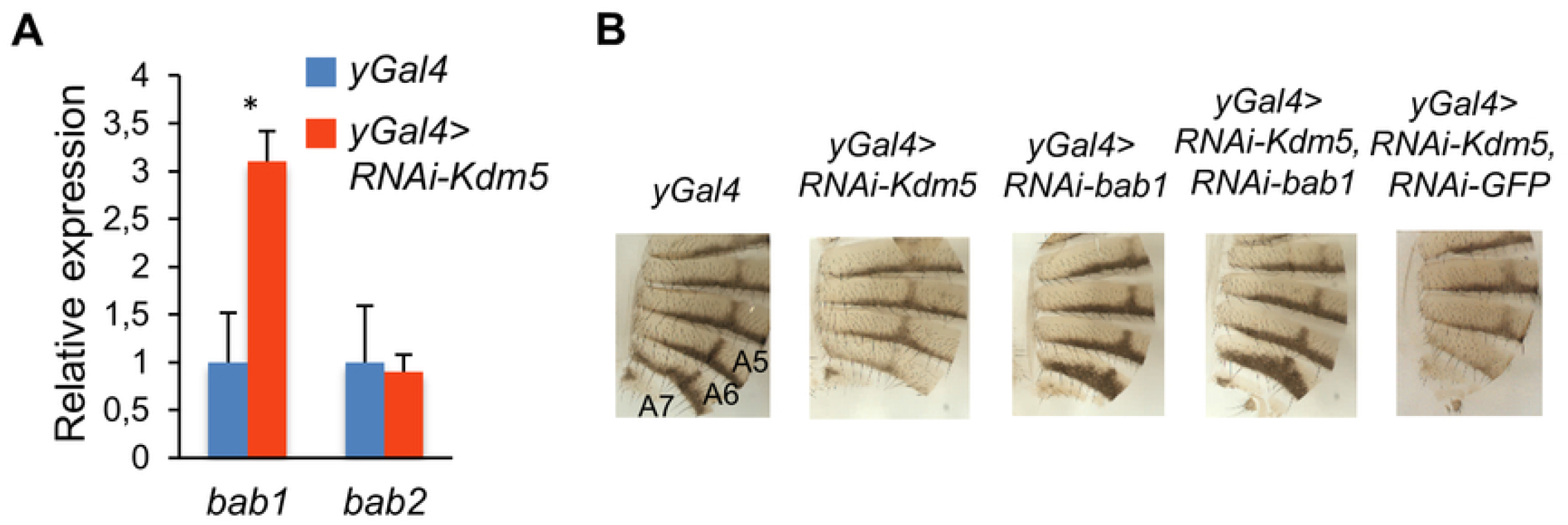
The activation of *tan* expression by Kdm5 is mediated, at least partly, by Bab1, a repressor of *tan* that is repressed by Kdm5. A. Quantification of the expression of *bab1* and *bab2* in pupal posterior abdominal epidermes (A5, A6, A7 segments) from control females (*yGal4*) or females with a down-regulation of *Kdm5* (*yGal4>RNAi-Kdm5*). The expression levels in *yGal4>RNAi-Kdm5* flies were normalized to the expression levels in *yGal4* flies. The expression of *bab1* but not *bab2* was significantly increased by *Kdm5* down-regulation. Pools of 30 epidermes, n=3 for all measurements except for *bab1* in *yGal4>RNAi-Kdm5* (n=2), normalization using the geometric mean of *RP49* and *Spt6* reference genes, t-test: *: p<0.05. B. Abdominal cuticles of heterozygous females either control (*yGal4*), expressing a *RNAi-Kdm5* transgene, a *RNAi-bab1* transgene, these two transgenes together, or the *RNAi-Kdm5* transgene and a *RNAi-GFP* transgene. Flies with *Kdm5* down-regulation were less pigmented than control flies, whereas flies with *bab1* down-regulation were more pigmented than control flies. *yGal4>RNAi-Kdm5, RNAi-bab1* flies showed the same pigmentation increase as *yGal4>RNAi-bab1* flies showing that *bab1* is epistatic on *Kdm5*. This was not due to Gal4 titration, as *yGal4>RNAi-Kdm5, RNAi-GFP* flies presented the same phenotype as *yGal4>RNAi-Kdm5* flies.

Taken together these results show that, among pigmentation genes, *tan* is the main target of Kdm5, with Kdm5 activating its expression. Furthermore, the regulation of *tan* by Kdm5 is mediated at least partly by *bab1,* a repressor of *tan* whose expression is repressed by Kdm5.

### Kdm5 and Trx depletions affect the transcription of many common genes in the female pupal abdominal epidermis, including several genes involved in the piRNA pathway

As *Kdm5* and *trx* both control the level of the H3K4me3 mark and are involved in female abdominal pigmentation by at least the activation of *tan* in young adults, we wondered whether they regulate other common genes acting earlier in the pigmentation process. We thus carried out a transcriptomic analysis by RNA sequencing (RNA-seq) of the posterior abdominal epidermes of old female pupae from control flies (*yGal4*) or flies with a down-regulation of *Kdm5* or *trx* (*yGal4>RNAi-Kdm5* and *yGal4>RNAi-trx*, respectively). These analyses identified 380 genes deregulated by Trx depletion (202 down-regulated and 178 up-regulated genes) (Supplementary Table 7) and 724 genes deregulated by Kdm5 depletion (454 down-regulated and 270 up-regulated genes) (Supplementary Table 8). As expected from Figure 3A, *bab1* was present in the list of genes up-regulated by Kdm5 depletion (log2 fold-change: 1.35), but its expression was not altered by Trx depletion. The gene *deadhead* (*dhd*), previously identified as a direct target of Kdm5 [24], was also strongly down-regulated in our *Kdm5* transcriptome (log2 fold-change: -2.63). As in already published *Kdm5* transcriptomes [22,24,32,33,48], most of the deregulated genes were moderately affected (less than 4-times for 611 genes, *i.e* - 2<log2 fold-change<2). Comparison of our *Kdm5* transcriptome with those of wing imaginal discs from a null *Kdm5* mutant [22] or of ovaries from RNAi-*Kdm5* females [24] confirmed the tissue and developmental stage-specificity of the previously described Kdm5 targets [22,33] (Figure 4A-B, Supplementary Table 8). Indeed, only 9.5% (n=70) of the genes deregulated in the abdominal epidermis were also deregulated in wing imaginal discs (among which only 47 were deregulated in the same direction). Similarly, only 2.3% (n=17) of the genes deregulated in the abdominal epidermis were also deregulated in the ovary (among which only 14 were deregulated in the same direction). Only two genes, *CG10814* (encoding an enzyme involved in lysine biosynthesis) and *SLC5A11* (encoding a sodium/solute transporter) were deregulated in the three tissues.

**Figure 4.**
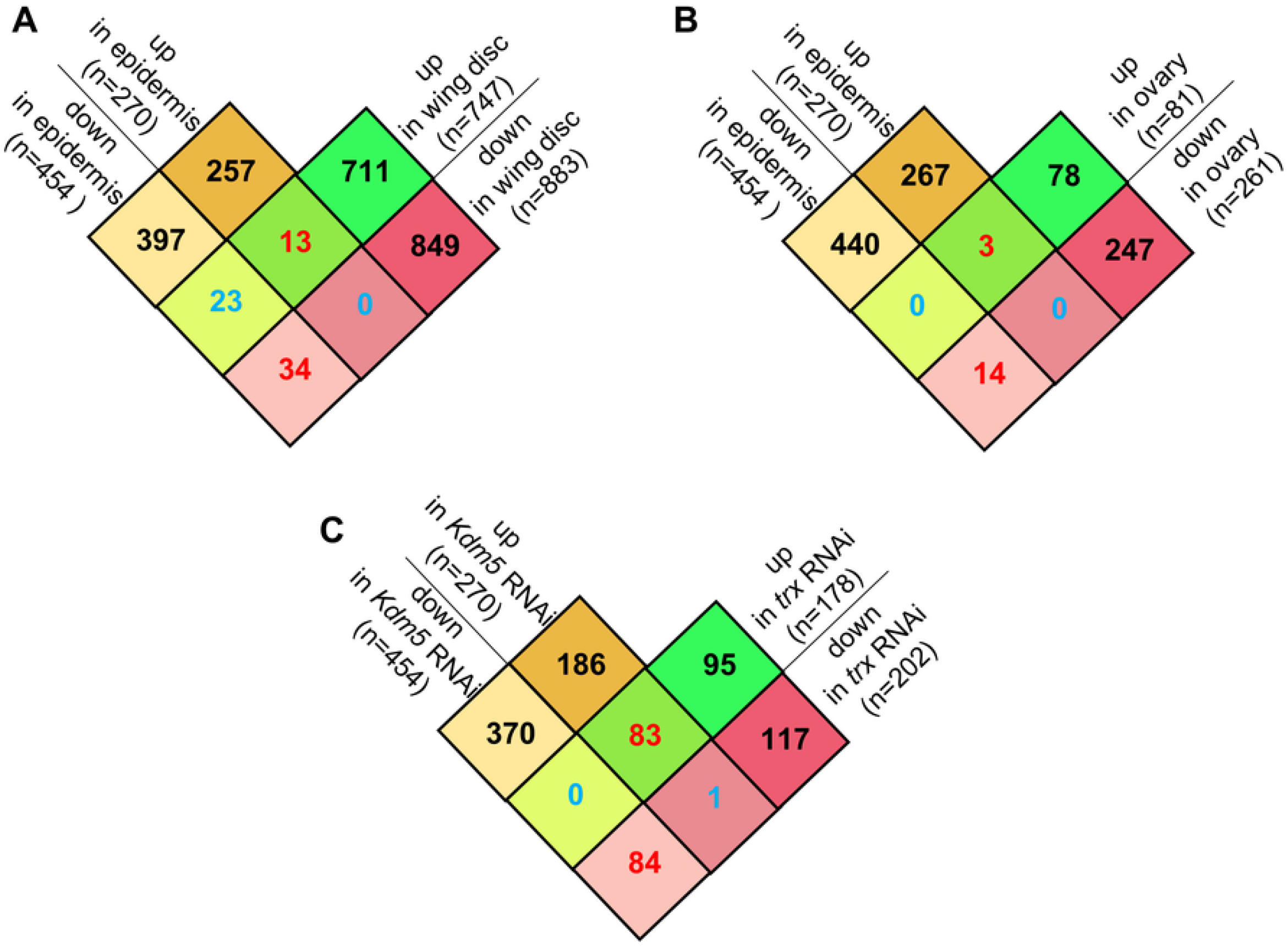
Comparison of genes deregulated by Kdm5 depletion in different tissues and with genes deregulated by Trx depletion. A, B. Venn diagrams showing comparison of the genes deregulated in our *Kdm5 RNAi* pupal abdominal epidermis transcriptome and in the *Kdm5* mutant wing disc transcriptome of Drelon *et al*. [22] (A) or in the *Kdm5 RNAi* ovary transcriptome of Torres-Campana *et al*. [24] (B). C. Venn-diagram showing comparison of genes deregulated in our *Kdm5-RNAi* and *trx-RNAi* pupal abdominal epidermis transcriptomes. Up: up-regulated genes; Down: down-regulated genes. Black numbers: genes not shared by the compared transcriptomes. Red numbers: genes deregulated in the same direction in the compared transcriptomes. Blue numbers: genes deregulated in opposite directions in the compared transcriptomes. Among the 724 genes deregulated in our *Kdm5* transcriptome, only 70 and 17 were also deregulated in the wing disc and ovary transcriptomes, respectively. In contrast, 168 of the genes deregulated in the *Kdm5* transcriptome were also deregulated in the *trx* transcriptome, mainly in the same direction.

Interestingly, comparison of the *Kdm5* and *trx* transcriptomes revealed a high degree of overlap for the deregulated genes (Figure 4C, Supplementary Table 9). Indeed, 23.2% (n=168) of the genes deregulated in the *Kdm5* transcriptome were also deregulated in the *trx* transcriptome, mainly in the same direction (83 genes and 84 genes up- or down-regulated in both conditions, respectively; only 1 gene deregulated in the opposite direction). These results suggest that *Kdm5* and *trx* act coordinately to regulate several genes in the female pupal abdominal epidermis.

Gene ontology (GO) analyses of the genes deregulated by Kdm5 or Trx depletion revealed various significantly enriched terms (Tables 1 and 2, adjusted p-value<0.05). Interestingly, several genes down-regulated by Kdm5 depletion were involved in piRNA metabolism (categories “P granule”, “Yb body”, “piRNA metabolic process”, “gene silencing by RNA”, “negative regulation of transposition, RNA mediated”), and some of them (*aub*, *del*, *SoYb*, *spn-E*) were also down-regulated by Trx depletion (Supplementary Table 9). It was also the case for *piwi*, significantly deregulated in the *Kdm5* transcriptome (log2 fold-change: -2.15; p: 0.00034) and with a p-value in the *trx* transcriptome just above significance (log2 fold-change: -1.45; p: 0.056). For *aub*, *del*, piwi, *SoYb* and *spn-E*, RNA-seq results were confirmed by RT-qPCR experiments on the posterior abdominal epidermis of female pupae (Supplementary Figure 4).

**Table 1.** Enriched gene ontology (GO) categories for genes down-regulated in the *RNAi-Kdm5* pupal epidermis transcriptome. For up-regulated genes, no category was significantly enriched.

| *GO term* | *Count* | *p-value* | *Benjamini adjusted p-value* |
| --- | --- | --- | --- |
| CC-GO:0043186~P granule | 11 | 5.35E-7 | 1.20E-4 |
| CC-GO:0070725~Yb body | 5 | 1.98E-5 | 2.23E-3 |
| CC-GO:0005777~peroxisome | 12 | 8.99E-5 | 6.74E-3 |
| CC-GO:0005874~microtubule | 13 | 3.07E-4 | 1.55E-2 |
| CC-GO:0005887~integral component of plasma membrane | 28 | 3.45E-4 | 1.55E-2 |
| CC-GO:0016020~membrane | 67 | 1.05E-3 | 3.96E-2 |
| BP-GO:0030717~karyosome formation | 9 | 8.82E-7 | 7.55E-4 |
| BP-GO:0034587~piRNA metabolic process | 8 | 5.36E-6 | 2.29E-3 |
| BP-GO:0031047~gene silencing by RNA | 7 | 1.88E-5 | 5.37E-3 |
| BP-GO:0055085~transmembrane transport | 26 | 3.78E-5 | 6.98E-3 |
| BP-GO:0048477~oogenesis | 20 | 4.07E-5 | 6.97E-3 |
| BP-GO:0045739~positive regulation of DNA repair | 4 | 7.59E-5 | 1.08E-2 |
| BP-GO:0008298~intracellular mRNA localization | 6 | 1.52E-4 | 1.86E-2 |
| BP-GO:0010526~negative regulation of transposition, RNA-mediated | 4 | 3.64E-4 | 3.82E-2 |
| BP-GO:0007052~mitotic spindle organization | 8 | 4.02E-4 | 3.83E-2 |
| BP-GO:0007315~pole plasm assembly | 5 | 5.57E-4 | 4.78E-2 |
| BP-GO:0046011~regulation of oskar mRNA translation | 4 | 6.25E-4 | 4.86E-2 |

**Table 2:** Enriched gene ontology (GO) categories for genes down-regulated (D) or up-regulated (U) in the *RNAi-trx* pupal epidermis transcriptome.

| *Deregulation* | *GO term* | *Count* | *p-value* | *Benjamini adjusted p-value* |
| --- | --- | --- | --- | --- |
| D | BP-GO:0055085~transmembrane transport | 20 | 2.89E-7 | 1.82E-4 |
| D | BP-GO:0046011~regulation of oskar mRNA translation | 4 | 6.56E-5 | 2.06E-2 |
| D | MF-GO:0022857~transmembrane transporter activity | 17 | 2.24E-8 | 5.29E-6 |
| U | CC-GO:0005739~mitochondrion | 25 | 5.53E-7 | 5.42E-5 |
| U | CC-GO:0005789~endoplasmic reticulum membrane | 13 | 9.74E-4 | 4.77E-2 |
| U | BP-GO:0006635~fatty acid beta-oxidation | 6 | 1.89E-5 | 5.79E-3 |
| U | BP-GO:0046680~response to DDT | 4 | 1.94E-4 | 2.97E-2 |

We thus wondered whether the piRNA pathway or some of its actors participate in the establishment of female abdominal pigmentation. We focused on the three most expressed candidate genes in the pupal abdominal epidermis: (1) *aubergine* (*aub*) (mean reads: 155.01; log2 fold-change in the *Kdm5* transcriptome: -1.51); (2) *piwi* (mean reads: 179.04; log2 fold-change in the *Kdm5* transcriptome: -2.15); (3) *spindle-E* (*spn-E*) (mean reads: 122.3; log2 fold-change in the *Kdm5* transcriptome: -1.73). *Piwi* and *aub* encode ARGONAUTE proteins of the PIWI family. In the ovary, Piwi and Aub associate with piRNA to form the pi-RISC complexes responsible for transcriptional and post-transcriptional repression of transposable elements (for a review, [49]). *spn-E* encodes an RNA-helicase. In ovarian germ cells, Spindle-E is involved in the ping-pong mechanism allowing primary piRNA amplification [50].

### Piwi and Aubergine act in abdominal epidermis, without evident piRNA production, to regulate female abdominal pigmentation through the regulation of *tan*

To test the impact of Aub, Piwi, or Spn-E depletion on female abdominal pigmentation, we analyzed females in which the tested gene was down-regulated by RNAi using the *yGal4* driver. These flies were compared with *yGal4* heterozygous flies (Figure 5A-B) and with heterozygous *RNAi* flies without the driver (Figure 5C-D). Compared to the *yGal4* and *RNAi* controls, Aub, Piwi and Spn-E depletions induced a significant decrease of A6 pigmentation. A5 depigmentation was observed for Aub with the two controls and for Piwi only with the *yGal4* control. A7 was depigmented only for Aub and Piwi, and only with the *yGal4* control. To confirm these phenotypic effects, we measured the expression of several pigmentation genes in the abdominal epidermis (segments A5 to A7) of *yGal4>RNAi-aub* and *yGal4>RNAi-piwi* females, as down-regulation of these two genes induced the strongest effect on pigmentation. *tan* was strongly deregulated by Aub and Piwi depletion (fold-change: 0.12; p<0.05). *ebony* was also down-regulated by Aub depletion, although to a lesser extent (fold-change: 0.65; p<0.01) (Figure 5E). In conclusion, these results demonstrate that down-regulating *aub* or *Piwi* in the abdominal epidermis decreases female abdominal pigmentation in a manner similar to down-regulation of *Kdm5*, and that these three genes participate in *tan* activation. As *aub* and *piwi* are both activated by Kdm5, they could act downstream of Kdm5 to regulate *tan* expression.

**Figure 5.**
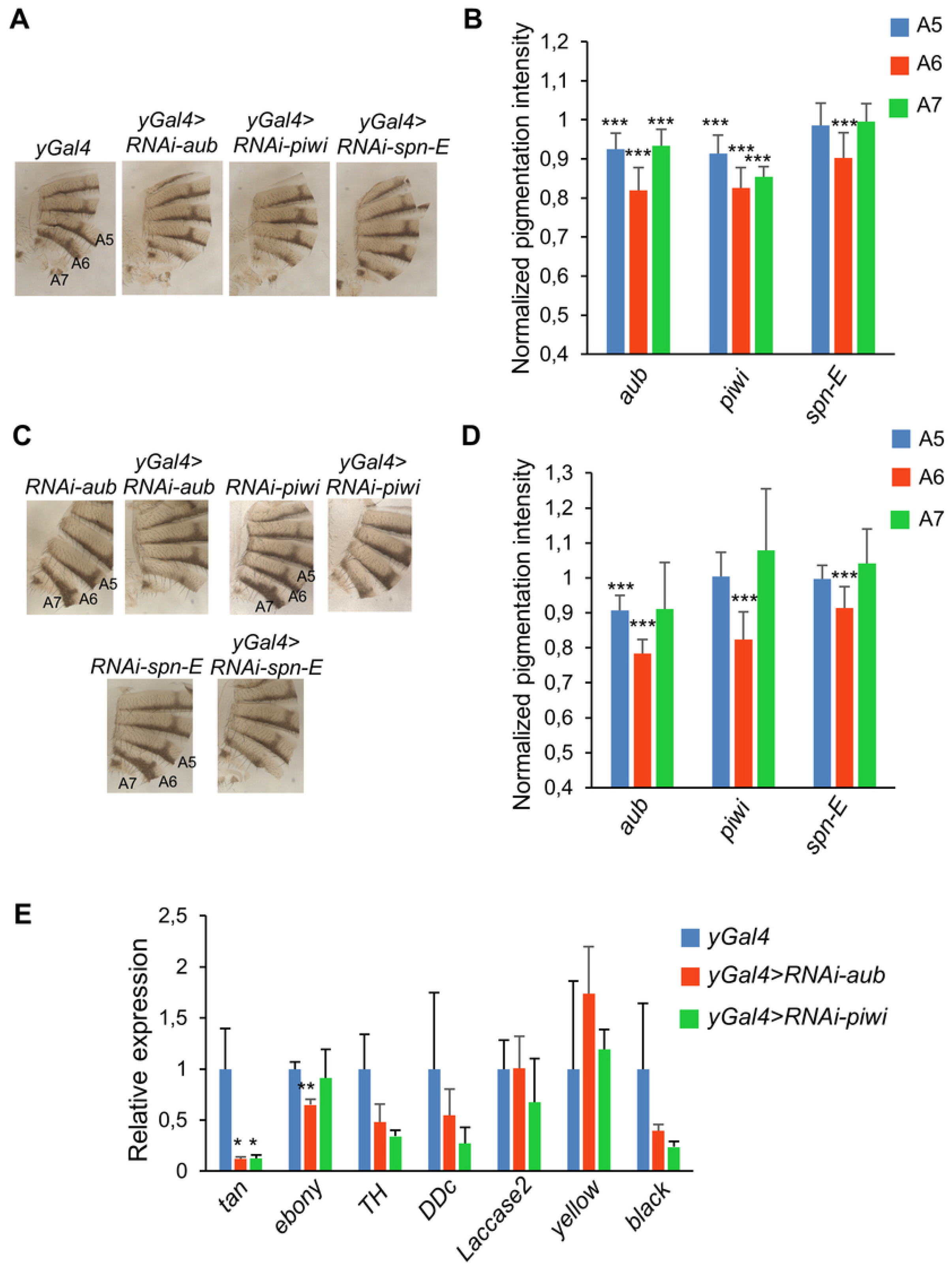
Effect of Aub, Piwi and Spn-E depletions on female abdominal pigmentation and pigmentation gene expressions. A, B. Abdominal cuticles (A) and quantification of A5, A6, A7 pigmentation (B) in control females (*yGal4*) or females expressing a RNAi transgene targeting *aub*, *piwi* or *spn-E* driven by *yGal4* (*yGal4>RNAi*). C, D. Abdominal cuticles (C) and quantification of A5, A6, A7 pigmentation (D) in control females (*RNAi*) or females expressing a RNAi transgene targeting *aub*, *piwi* or *spn-E* driven by *yGal4* (*yGal4>RNAi*). In B and D, pigmentation intensities of the *yGal4>RNAi* flies were normalized to pigmentation intensities of the control flies (n=15 *per* genotype, t-test; ***: p<0.001). Compared to *yGal4* control flies (B), down-regulation of *aub* and *piwi* induced a decrease of A5, A6 and A7 pigmentation whereas depigmentation was observed only in A6 for *spn-E* down-regulation. Compared to *RNAi* control flies (D), a pigmentation decrease was observed in A5 and A6 for *yGal4>RNAi-aub* flies, and only in A6 for *yGal4>RNAi-piwi* and *y-Gal4>RNAi-spnE* flies. E. Quantification of pigmentation gene expressions in posterior abdominal epidermes (A5, A6, A7 segments) of heterozygous young females either control (*yGal4*), or with a down-regulation of *aub* or *piwi* (*yGal4>RNAi-aub* and *yGal4>RNAi-piwi*, respectively). The expression levels in *yGal4>RNAi* flies were normalized to the expression levels in *yGal4* flies. The level of *tan* expression was strongly lowered by *aub and piwi* down-regulation (pools of 30 epidermes, n=3, normalization using the geometric mean of *RP49* and *Spt6* reference genes, t-test: *: p<0.05; **: p<0.01).

To detect potential piRNA in the female pupal abdominal epidermis, we performed small RNA sequencing experiments (small-RNAseq) and analyzed the quantity, size distribution and sequence composition of the 19-29-nt RNA. We compared heterozygous *yGal4* control flies to *yGal4>RNAi-Kdm5* and *yGal4>RNAi-piwi* flies. To rule out the possibility of a contamination of pupal abdominal epidermes by ovarian tissue during dissection, small-RNAseq was also performed on abdominal epidermes of homozygous *ovoD1* females that exhibited degenerated ovaries [51]. We then compared these abdominal epidermis small-RNAseq data with small-RNAseq data from adult ovary of control females (GEO accession number GSE203279, [52]). For comparison, the number of reads in each library was normalized on one million (rpm). After computational removal of annotated small RNA (*i.e.* t-r-mi-sno-RNA), the quantity of 19-29-nt reads, called thereafter cleaned reads, was lower in the abdominal epidermis libraries (16.10^4^ to 27.10^4^) than in the ovary library (5.10^5^). As expected, in the *yGal4>RNAi-Kdm5* and *yGal4>RNAi-piwi* libraries, a high proportion of the cleaned reads were 21-22-nt siRNA against *Kdm5* or *piwi* (19% and 18% of the cleaned reads, respectively) (Supplementary Table 5 and Supplementary Figure 5A). The 23-29-nt category, corresponding to potential piRNA, was much more abundant in the ovary library (46.10^4^) than in the abdominal epidermis libraries (between 14.10^3^ and 34.10^3^ depending on the genotype) (Supplementary Table 5 and Figure 6A). Several observations suggested that the majority of 23-29-nt RNA in abdominal epidermis libraries were probably not piRNA: (1) they did not show a canonical piRNA size distribution (Figure 6B, to be compared with the ovary library size distribution in Figure 6A); (2) the ovarian 23-29-nt RNA exhibited a strong bias for a uracil in first position (f=0.71), which is a canonical piRNA signature. In contrast, this bias did not exist for the abdominal epidermis 23-29-nt RNA (f between 0.28 and 0.46) (Figure 6C); (3) a very low percentage of the 23-29-nt unique reads in abdominal epidermis libraries mapped on the major piRNA cluster loci from which piRNA are produced [53] (0.4 to 6% depending on the genotype), whereas this percentage was 61.4% in the ovary library (Supplementary Table 5). In conclusion, these results reveal that piRNA are absent or very rare in the pupal abdominal epidermis.

**Figure 6.**
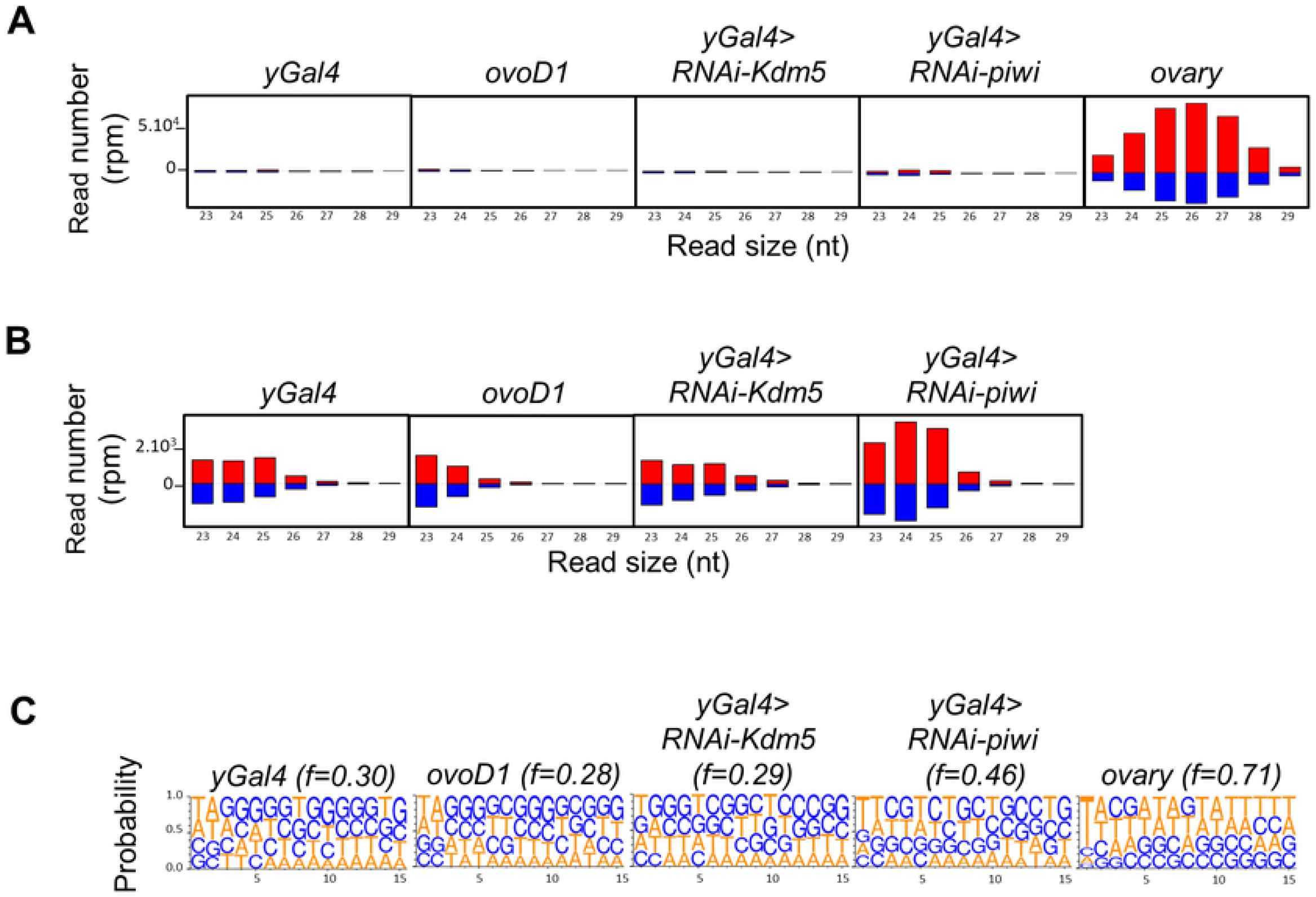
piRNA are absent or very rare in pupal abdominal epidermis. A. Size distribution (in reads per million, rpm) of the 23-29-nt RNA from pupal abdominal epidermis libraries (*yGal4*, *ovoD1*, *yGal4>RNAi-Kdm5*, *yGal4>RNAi-piwi* females) and adult ovary library, after removal of t-RNA, mi-RNA, r-RNA and other annotated small RNA. Red and blue bars correspond to sense and antisense reads, respectively. In the abdominal epidermis libraries, very few 23-29-nt RNA, corresponding to potential piRNA, were present. By contrast, they were very abundant in the ovary library, as expected, and presented a size distribution typical of piRNA. C. Enlarged view of the size distribution of 23-29-nt RNA from abdominal epidermis libraries, showing no typical piRNA size distribution. C. Logo graphs showing sequence composition of the 23-29-nt RNA. In the ovary, 23-29-nt RNA had a high tendency to begin with a uracil (T in the graph, f=0.71), a typical piRNA signature, whereas it was not the case for 23-29-nt RNA from pupal abdominal epidermis.

### Kdm5 and Piwi play a role in the pupal fat body to regulate female abdominal pigmentation despite no piRNA detection

Recently, Piwi was shown to be expressed in the larval fat body, although the existence of piRNA in this tissue was not investigated [54]. In the larval fat body, Piwi depletion induced over-expression of ImpL2, an IGF-binding protein that negatively regulates the insulin/insulin-like signaling pathway (IIS) [55]. This result caught our attention because the IIS is required for proper body pigmentation. Indeed, RNAi down-regulation in the female abdominal epidermis of *InR*, which encodes the insulin receptor, induces a strong abdominal depigmentation [56]. We thus wondered whether, during the pupal stage, Piwi, independently or not of piRNA, could act also in the fat body cells to regulate abdominal pigmentation. In the pupa, during metamorphosis, the larval fat body undergoes remodeling by tissue dissociation, giving rise to isolated or clumps of fat body cells that persist in the adult abdomen until a few days after eclosion [57]. RT-qPCR experiments revealed that the expression of *piwi* previously detected in the larval fat body persisted during the pupal stage (Figure 7A). As in the pupal epidermis, *piwi* expression in the pupal fat body was much lower than in the adult ovary, whereas not significantly different from its expression in the pupal abdominal epidermis (fold-change between ovary and fat body: 0.056; fold-change between ovary and epidermis: 0.17).

**Figure 7.**
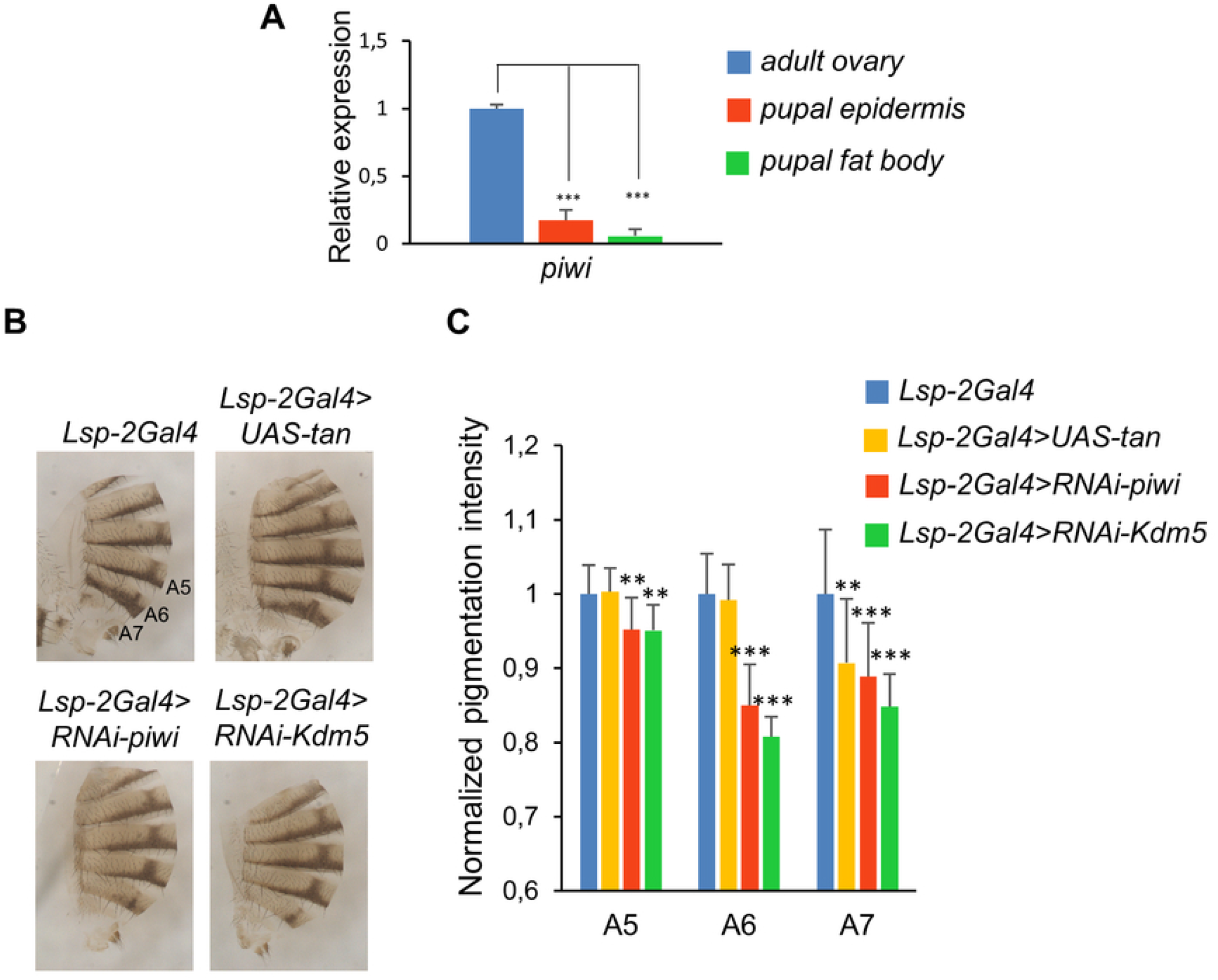
Piwi and Kdm5 depletions in the fat body induce a decrease of female abdominal pigmentation. A. Quantification of *piwi* expression in the pupal fat body of heterozygous *Lsp-2Gal4* females, compared to its expression in wild-type adult ovaries and in the pupal abdominal epidermis from heterozygous *yGal4* females. The expression levels in the pupal fat body and abdominal epidermis were normalized to the expression level in ovary (3 replicates *per* tissue, normalization using the geometric mean of *RP49* and *Spt6* reference genes, t-test: ***: p<0.001). *piwi* expression in the pupal fat body and abdominal epidermis was much lower than in ovary. B, C. Abdominal cuticles (B) and quantification of A5, A6, A7 pigmentation (C) in control females (*Lsp-2Gal4*) or females expressing, under the control of *Lsp-2Gal4*, a *UAS-tan* transgene or a RNAi transgene targeting *piwi* or *Kdm5*. In C, pigmentation intensities of the *Lsp-2Gal4* flies were used for normalization (n=15 *per* genotype, t-test; **: p<0.01; ***: p<0.001). The over-expression of *tan* in the fat body did not increase pigmentation. In contrast, down-regulation of *piwi and Kdm5* in the fat body induced a decrease of A5, A6 and A7 pigmentation.

To test whether Piwi could act in the fat body to regulate abdominal pigmentation, its expression was down-regulated by RNAi using the *Lsp-2Gal4* driver. The *Larval serum protein-2* gene (*Lsp-2*) encodes an hemolymph protein specifically produced in the fat body from the larval stage to the adulthood [58–60]. We first checked, using a *UAS-H2B-YFP* reporter transgene, that the *Lsp-2Gal4* driver was only expressed in the larval and pupal fat body and not in the pupal abdominal epidermis (Supplementary Figure 6A). On the opposite, the *yGal4* driver, not expressed at the larval stage, was excluded from the pupal fat body and restricted to abdominal epidermis (Supplementary Figure 6B). Over-expressing *tan* using the *Lsp-2Gal4* did not increase abdominal pigmentation, thus confirming that the *Lsp2-Gal4* driver was not expressed in the abdominal epidermis (Figure 7B-C). Indeed, as shown in Figure 2B and previously described [13,47], *tan* over-expression in the abdominal epidermis increased pigmentation. Interestingly, Piwi and Kdm5 depletion in the pupal fat body induced a decrease of A5, A6 and A7 pigmentation (Figure 7B-C). Moreover, opposite to what was observed in the abdominal epidermis, *piwi* expression in the pupal fat body was not regulated by Kdm5 (Supplementary Figure 7A).

We then wondered whether, during the pupal stage, Piwi could participate in the control of the IIS pathway through repression of its negative regulator ImpL2, as was shown in the larval fat body [54]. Down-regulation of *piwi* in the pupal abdominal epidermis (*yGal4* driver) or the pupal fat body (*Lsp-2Gal4* driver) did not modify *ImpL2* expression (S7B Fig). Moreover, no important change in pigmentation was observed when *ImpL2* was down-regulated in the pupal abdominal epidermis or the pupal fat body) (Supplementary Figure 7C-D). By contrast, a strong decrease of pigmentation was observed when *InR*, encoding the insulin receptor, was down-regulated in the two tissues (Supplementary Figure 7E-F). The depigmentation of *ygal4>RNAi-InR* flies confirmed that, as previously shown [56], IIS acts in a cell autonomous manner in the abdominal epidermis to regulate melanin production. As the same effect was induced using the *Lsp2-Gal4* driver, it demonstrates that regulation of the IIS in the pupal fat body exerts also a systemic effect on melanin production.

To question the presence of piRNA in pupal fat body, we sequenced the 19-29-nt small RNA from the pupal fat body of *Lsp-2Gal4*, *Lsp-2Gal4>RNAi-Kdm5* and *Lsp-2Gal4>RNAi-piwi* flies. As previously, we normalized the read numbers of each library to one million (rpm). The quantity of 23-29-nt reads in pupal fat body libraries after computational removal of annotated small RNA, called thereafter cleaned reads, was very low (56.10^4^ to 11.10^4^). As expected, in the *Lsp-2Gal4>RNAi-Kdm5* and *Lsp2-Gal4>RNAi-piwi* libraries, a high proportion of the cleaned reads were 21-22-nt siRNA against *Kdm5* or *piwi* (5.7% and 19.7% of the cleaned reads, respectively) (Supplementary Table 6 and Supplementary Figure 5B). As in the abdominal epidermis libraries, the 23-29-nt RNA, corresponding to potential piRNA, were very scarce (between 15.10^4^ and 36. 10^4^ depending on the genotype) (Supplementary Table 6 and Figure 8A) and were not reduced by Piwi depletion. These 23-29-nt reads exhibited neither the typical piRNA size distribution (Figure 8B), nor a bias for a uracil in first position (f between 0.14 and 0.20) (Figure 8C). Lastly, a very low percentage of 23-29-nt unique reads mapped on the piRNA cluster loci (0.4 to 3.6%) (Supplementary Table 6). We thus concluded that, as in the abdominal epidermis, piRNA are absent or very rare in the pupal fat body.

**Figure 8.**
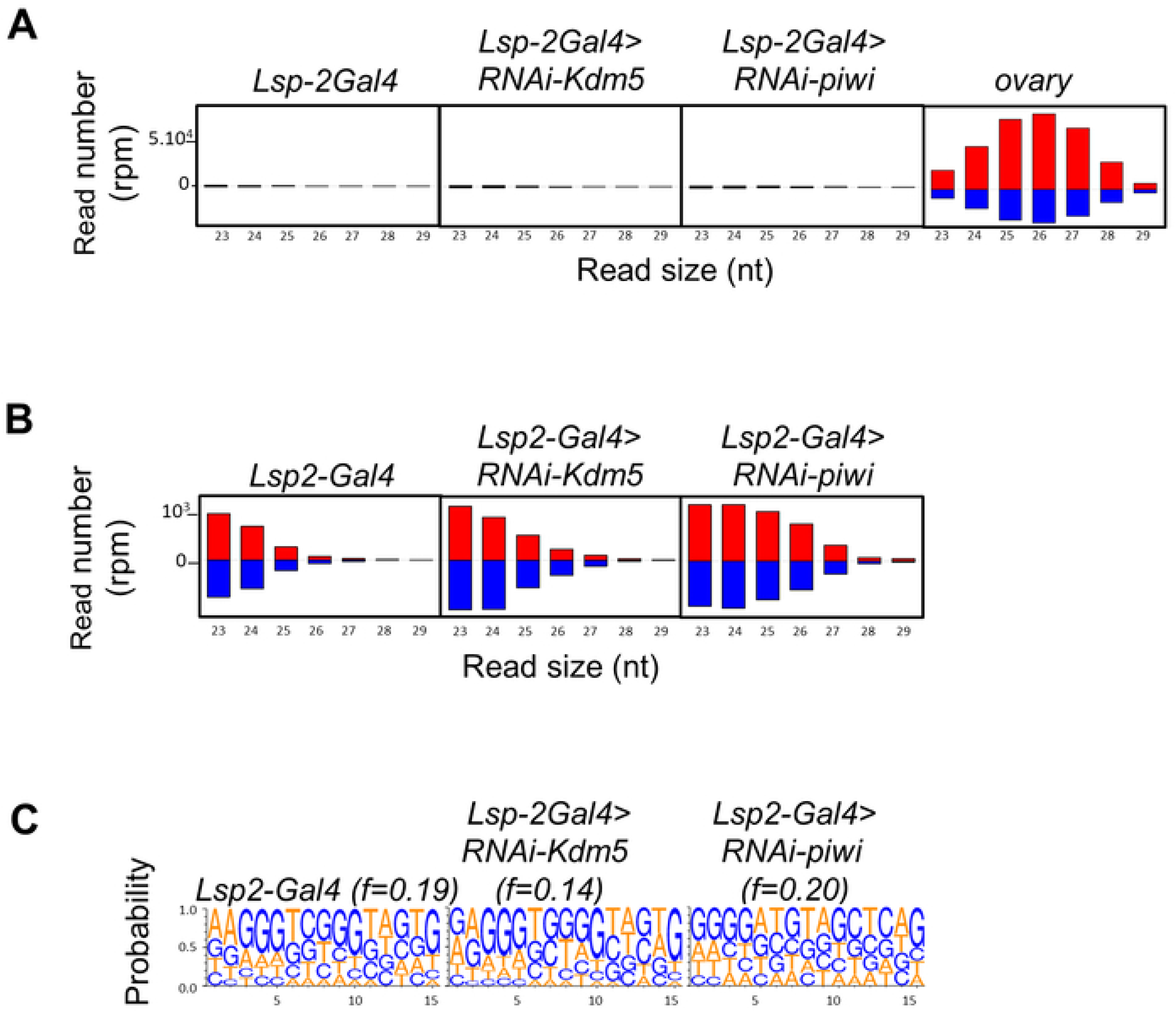
piRNA are absent or very rare in the pupal fat body. A. Size distribution (in reads per million, rpm) of the 23-29-nt RNA from pupal fat body after removal of t-RNA, mi-RNA, r-RNA and other annotated small RNA. Red and blue bars correspond to sense and antisense reads, respectively. 23-29-nt RNA, corresponding to potential piRNA, were much less abundant in pupal fat body libraries than in ovary library. B. Enlarged view of the size distribution of 23-29-nt RNA from pupal fat body libraries showing no typical piRNA size distribution. C. Logo graphs showing sequence composition of the 23-29-nt RNA from pupal fat body library. They did not present a bias for a uracil in first position (T in the graphs), a typical piRNA signature.

In conclusion, our results show that, during the pupal stage, *Kdm5* and *piwi* act in two different tissues, the fat body and the abdominal epidermis, to regulate female abdominal pigmentation. In the fat body, *Kdm5* and *piwi* may act independently to modulate one or several pathways exerting a systemic effect on pupal abdominal epidermis. In the pupal abdominal epidermis, *Kdm5* activates the expression of the pigmentation gene *tan* through at least the regulation of *piwi*, *bab1* and *aub*, although we could not exclude a role of *Kdm5* independently of these three genes. As we were not able to identify piRNA in pupal abdominal epidermis and pupal fat body, it is likely that *piwi* achieves its function in these tissues without any production of piRNA.

## Discussion

In this study, we have shown that the histone demethylase Kdm5, erasing the H3K4me3 mark, participates during pupal life in the establishment of female abdominal pigmentation. Kdm5 exerts its effect through transcriptional activation of the pigmentation gene *tan* in the abdominal epidermis of young adult females. Analysis of *tan-GFP* transgene expression revealed that Kdm5 does not regulate the activity of the *t-MSE* abdominal epidermis enhancer but acts more likely on the promoter of *tan*. Interestingly, our previous studies revealed that *tan* was also activated by the histone-methyl transferase Trx and that the level of H3K4me3 on the promoter of *tan* decreased when *trx* was down-regulated [13]. Thus, *tan* could be a direct target of both Trx and Kdm5, regulating together the level of H3K4me3 on its promoter according to the mechanism proposed in [34]. In addition to this potential direct effect, Kdm5, alone or with Trx, may also directly or indirectly regulate some genes acting earlier during development and regulating the expression of *tan*. Indeed, our transcriptomic analyses on pupal abdominal epidermes, showing a huge overlap between genes deregulated by Kdm5 or Trx depletion, suggest that these two proteins act coordinately to regulate several genes in this tissue and that regulation of these common targets requires the demethylase activity of Kdm5. Further experiments, such as chromatin immunoprecipitations, are required to identify the direct targets of Kdm5 and Trx in the pupal abdominal epidermis. As for the genes deregulated only in the *Kdm5* transcriptome, such as *bab1*, encoding a repressor of *tan* [8], Kdm5 could act on their H3K4me3 levels in cooperation with other chromatin factors required for H3K4 methylation. One candidate could be Ash2, previously shown to directly regulate several direct targets of Kdm5 [34]. Alternatively, the effect of Kdm5 on these genes could be either indirect or independent of its demethylase activity. Comparison of abdominal epidermis transcriptomes of wild-type flies and flies expressing, as the only source of Kdm5, a mutated form lacking demethylase activity (point mutation in the JmjC domain, [22,23]) would allow to answer this question. As for the importance of the demethylase activity of Kdm5 in female abdominal pigmentation establishment, comparison of abdominal pigmentation of wild-type females and females expressing a Kdm5 mutated form lacking demethylase activity did not allow us to conclude, probably because of the different genetic backgrounds in the tested lines. It is also possible that Kdm2, another histone demethylase erasing H3K4me3 [61,62] acts redundantly to Kdm5 in the process of abdominal pigmentation as this is the case during larval development. Indeed, developmental lethality of *Kdm2,Kdm5* double mutants is much stronger than that of either single mutant. Moreover, this lethality is rescued only by a wild-type form of Kdm5 and not by a form defective in demethylase activity, suggesting that Kdm5 and Kdm2 act redundantly in the regulation of H3K4me3 to ensure proper development [21].

Among the transcriptional targets of Kdm5 in the pupal abdominal epidermis, we identified several genes involved in the piRNA pathway, such as *aub*, *piwi*, *SoYB*, *spn-E*. This could reveal the existence of a coordinated transcription of these genes in the pupal abdominal epidermis, as shown in ovaries with the transcription factor Ovo identified as a direct regulator of several piRNA pathway factors [63]. We demonstrated a new somatic function of the ARGONAUTE proteins Piwi and Aub in the pupal abdominal epidermis, acting downstream of Kdm5 and involved in the regulation of abdominal pigmentation via *tan* expression. Interestingly, down-regulation of *Kdm5*, *piwi* and *aub* not only in the pupal abdominal epidermis, but also in the pupal fat body, had an impact on abdominal pigmentation, thus revealing for the first time a role of the pupal fat body in the establishment of abdominal pigmentation. Insulin signaling could be involved in this process, as we showed that depletion of the insulin receptor in the pupal fat body induced a pigmentation decrease. However, we could not establish a functional link between Kdm5, Piwi, and the insulin signaling regulation, as previously shown in the larval fat body where Piwi depletion induced the over-expression of ImpL2, a negative regulator of insulin signaling [54]. A *RNAi* genetic screen with the fat body-specific driver *Lsp-2Gal4* could allow identification of the pathways mediating the effect of the fat body on abdominal pigmentation.

Although our study demonstrates the involvement of Kdm5, Piwi and Aub in the establishment of abdominal pigmentation through their action in two different pupal tissues, the molecular mechanisms by which they exert their effect remain to be investigated. It seems that Kdm5 and Piwi do not act similarly in the pupal abdominal epidermis and the pupal fat body. Indeed, in the abdominal epidermis, Kdm5 activates the expression of *piwi*, whereas it is not the case in the fat body. Our results suggest that the role of Piwi and Aub in abdominal pigmentation regulation could be piRNA-independent, as we could not detect *bona fide* piRNA in either tissue. However, we cannot exclude that the piRNA amount was too low to be detected. This question could be answered in future studies by generating oxidized small RNA libraries. Indeed, activated piRNA in pi-RISC complexes are 2’-O-methylated and can be enriched using sodium periodate oxidation [64]. It was shown recently that, in the drosophila adult intestine, oxidized small RNA libraries allowed to detect a very low amount of piRNA [40,41]. Similarly, in the adult fat body, piRNA were identified from oxidized libraries [38]. Finally, mass spectrometry analysis in the pupal fat body and the abdominal epidermis, as performed in early embryos [65], would allow identification of Piwi and Aub interactors in these tissues and thus a better understanding of their mode of action.

## Acknowledgments

We thank Valérie Ribeiro for her help in fly husbandry, Antoine Boivin for advice in small RNA-seq analyses and critical reading of the manuscript, the members of the team for stimulating discussions, Naïra Nouar from the InforBio Bioinformatics platform (IBPS) for the training of Raphael Narbey in RNA-seq analyses, the Bloomington Drosophila Stock Center and the Vienna Drosophila Resource Center for fly stocks.

## Supporting information

**Supplementary Figure 1. Quantification of the *RNAi-Kdm5* and *UAS-Kdm5* transgene efficiencies using the ubiquitous *daughterless-Gal4* (*daGal4*) driver.** RT-qPCR quantification of *Kdm5* expression in *daGal4>RNAi-Kdm5* larvae (A) or in *daGal4>UAS-Kdm5* embryos (B). Compared to *daGal4* individuals, the *RNAi-Kdm5* transgene induced a 4.5-fold down-regulation and the *UAS-Kdm5* transgene a 1.9-fold up-regulation (n=2 for *daGal4* and *daGal4>RNAi-Kdm5* individuals; n=3 for *daGal4>UAS-Kdm5* individuals; normalization using the geometric mean of *RP49* and *Spt6* reference genes, t-test: *: p<0.05).

**Supplementary Figure 2. Confirmation of the effect of *Kdm5* down-regulation on abdominal pigmentation using a second control, a second driver and a second RNAi line.** A, B. Abdominal cuticles (A) and quantification (B) of A5, A6 and A7 pigmentation of control females (*RNAi-Kdm5,* BDSC_28944 line) or females expressing the *RNAi-Kdm5* transgene driven by *yGal4* or *pnGal4*. C, D. Abdominal cuticles (C) and quantification (D) of A5, A6 and A7 pigmentation of control females (*RNAi-Kdm5*, BDSC_36652 line) or females expressing the *RNAi-Kdm5* transgene driven by *yGal4* or *pnGal4* (n=15 *per* genotype, t-test: *: p<0.05; **: p<0.01; ***: p<0.001).

**Supplementary Figure 3. Expression of *tan-GFP* reporter transgenes upon *Kdm5* down- or up-regulation.** A to D. Effect of *Kdm5* down-regulation (A, B) or over-expression (C, D) on the activity of *tan* regulatory sequences (reporter transgene *5’tan-GFP*). A, C: GFP fluorescence in A5, A6 and A7 abdominal segments of young control females (*5’tan-GFP, yGal4*) or females expressing a *RNAi-Kdm5* transgene (*5’tan-GFP, yGal4>RNAi-Kdm5*) or a *UAS-Kdm5* transgene (*5’tan-GFP, yGal4>UAS-Kdm5*). B, D: Quantification of GFP positive nuclei in A5, A6 and A7. The expression of the *5’tan-GFP* reporter transgene was lowered in A6 and A7 by *Kdm5* down-regulation and increased in A6 and A7 by *Kdm5* over-expression (n=10 *per* genotype, t-test: *: p<0.05; **: p<0.01; ***: p<0.001). E, F. No effect of *Kdm5* down- regulation on the activity of the *t-MSE* enhancer (reporter transgene *t-MSE-GFP*). E. GFP fluorescence in A5, A6 and A7 abdominal segments of young control females (*t-MSE-GFP, yGal4*) or females expressing a *RNAi-Kdm5* transgene (*t-MSE-GFP, yGal4>RNAi-Kdm5*). F. Quantification of GFP positive nuclei in A5, A6 and A7 (n=10 per genotype).

**Supplementary Figure 4. Several piRNA pathway genes are down-regulated by depletion of Kdm5 or Trx in the female pupal abdominal epidermis.** RT-qPCR quantification of the expression of several genes involved in piRNA metabolism in the posterior abdominal epidermis (A5, A6, A7 segments) of female pupae either control (*yGal4*) or with a down-regulation of *Kdm5* (*yGal4>RNAi-Kdm5)* or *trx (yGal4>RNAi-trx).* The expression levels in *Kdm5* and *trx* RNAi flies were normalized to the expression levels in *yGal4* flies using the geometric mean of *RP49* and *Spt6* reference genes (pools of 30 epidermes, n=3, t-test: (*): 0.05<p<0.17; *: p<0.05).

**Supplementary Figure 5. The cleaned 19-29-nt RNA are much less abundant in pupal abdominal epidermis and pupal fat body libraries than in adult ovary library.** Size distribution (in reads per million, rpm) of the 19-29-nt RNA from pupal abdominal epidermis (A), adult ovary (A) and pupal fat body (B), after removal of t-RNA, mi-RNA, r-RNA and other annotated small RNA. Red and blue bars correspond to sense and antisense reads, respectively. In A and B, the white asterisks on the 21-22-nt categories indicate the si-RNA against *Kdm5* in the *RNAi-Kdm5* libraries and against *piwi* in the *RNAi-piwi* libraries.

**Supplementary Figure 6. Tissue-specificity of the *Lsp-2Gal4* and *yGal4* drivers.** A. Expression in the fat body (FB) and abdominal epidermis (AE) of a third instar larva (up) or old pupa (bottom) of a *UAS-H2B-YFP* transgene driven by *Lsp-2Gal4*. H2B-YFP was expressed in the fat body but not in the abdominal epidermis. B. Expression in the fat body (FB) and abdominal epidermis (AE) of a third instar larva (up) or old pupa (middle and bottom) of a *UAS-H2B-YFP* transgene driven by *yGal4.* H2B-YFP was not expressed during the larval stage, whereas in pupa it was expressed in the abdominal epidermis but not in the fat body. In A and B, the merge pictures were obtained by superimposition of the nuclear (Hoechst) and YFP stainings.

**Supplementary Figure 7. Expression of *piwi* and *ImpL2* in pupal tissues and pigmentation phenotype induced by down-regulation of *ImpL2* or *InR* in the pupal abdominal epidermis or the fat body.** A. No effect of *Kdm5* down-regulation in the fat body on *piwi* expression (*Lsp-2Gal4* driver). B. No effect of *piwi* down-regulation on *ImpL2* expression in the abdominal epidermis (*yGal4* driver) or in the fat body (*Lsp-2Gal4* driver) (A and B: 3 replicates *per* genotype, normalization using the geometric mean of *RP49* and *Spt6* reference genes). C to F. Abdominal cuticles (C, E) and quantification (D, F) of A5, A6 and A7 pigmentation of control females (*RNAi-ImpL2* or *RNAi-InR)* or females expressing the *RNAi-ImpL2 or RNAi-InR* transgene in the abdominal epidermis (*yGal4* driver) or the fat body (*Lsp-2Gal4* driver). Whereas ImpL2 depletion induced no change in pigmentation, InR depletion in the two tissues induced depigmentation. N=15 *per* genotype, t-test: **: p<0.01, ***: p<0.001.

**Supplementary Table 1. List of fly lines used in this study.**

**Supplementary Table 2. List of the RNAi lines tested in our screen for abdominal pigmentation modification.**

**Supplementary Table 3: List of primers used in this study.**

**Supplementary Table 4. Characteristics of the RNAseq libraries from pupal abdominal epidermis.**

**Supplementary Table 5. Characteristics of the small-RNAseq libraries from pupal abdominal epidermis and adult ovary.**

**Supplementary Table 6. Characteristics of the small-RNAseq libraries from pupal fat body.**

**Supplementary Table 7. List of genes deregulated in *yGal4>RNAi-trx* pupal abdominal epidermes.**

**Supplementary Table 8. List of genes deregulated in *yGal4>RNAi-Kdm5* pupal abdominal epidermes.**

**Supplementary Table 9. List of genes deregulated in common in *yGal4>RNAi-Kdm5* and *yGal4>RNAi-trx* pupal abdominal epidermis.**

